# Dietary Exposure to Antibiotic Residues Facilitates Metabolic Disorder by Altering the Gut Microbiota and Bile Acid Composition

**DOI:** 10.1101/2021.07.06.451284

**Authors:** Rou-An Chen, Wei-Kai Wu, Suraphan Panyod, Po-Yu Liu, Hsiao-Li Chuang, Yi-Hsun Chen, Qiang Lyu, Hsiu-Ching Hsu, Tzu-Lung Lin, Ting-Chin David Shen, Yu-Tang Yang, Hsin-Bai Zou, Huai-Syuan Huang, Yu-En Lin, Chieh-Chang Chen, Chi-Tang Ho, Hsin-Chih Lai, Ming-Shiang Wu, Cheng-Chih Hsu, Lee-Yan Sheen

**Author notes:** There authors contributed equally for the study (Rou-An Chen, Wei-Kai Wu). **Corresponding authors:** Lee-Yan Sheen, No. 1, Sec. 4, Roosevelt Rd., Taipei, Taiwan 10617, Cheng-Chih Hsu, No. 1, Sec. 4, Roosevelt Rd., Taipei, Taiwan 10617.

## Abstract

Antibiotics used as growth promoters in livestock and animal husbandry can be detected in animal-derived food. Epidemiological studies have implicated that exposure to these antibiotic residues in food may be associated to childhood obesity. Herein, the effect of exposure to residual dose of tylosin—an antibiotic growth promoter—on host metabolism and gut microbiota was explored *in vivo*. Theoretical maximal daily intake (TMDI) doses of tylosin were found to facilitate high-fat diet-induced obesity, induce insulin resistance, and perturb the composition of gut microbiota in mice. The obesity-related phenotypes were transferrable to germ-free recipient mice, indicating that the effects of TMDI dose of tylosin on obesity and insulin resistance occurred mainly via alteration of the gut microbiota. Tylosin TMDI exposure restricted to early life, which is the critical period of gut microbiota development, altered the abundance of specific bacteria related to host metabolic homeostasis later in life. Moreover, early-life exposure to tylosin TMDI was sufficient to modify the ratio of primary to secondary bile acids, thereby inducing lasting metabolic consequences via the downstream FGF15 signaling pathway. Altogether, these findings demonstrate that exposure to very low dose of antibiotic residues, whether continuously or in early life, can exert long-lasting effects on host metabolism by altering gut microbiota and its metabolites.

**Importance:** Evidence has indicated that chronic exposure to antibiotic residues in food could contribute to obesity. However, few studies have investigated the effect of chronic exposure to very low-dose antibiotic residue in food (~1000-fold lower than the therapeutic dose) on gut microbiota and host metabolism. Our study demonstrates that even with limited exposure in early life, a residual dose of tylosin causes lasting metabolic disturbances through altering gut microbiota and its metabolites. Our findings reveal that the gut microbiota is susceptible to previously ignored environmental factors.

## Introduction

Antibiotics administrated at sub-therapeutic doses have been used as growth promoters since 1940s (1–3). According to the U.S. Food and Drug Administration, approximately two-thirds of all antimicrobial agents used in the United States are for livestock and animal husbandry, driven by the demand to improve the production of animal-derived foods (4). Antibiotics used in livestock may remain in animal-derived foods and contribute to inadvertent antibiotic residue consumption by humans. To avoid potential health hazards to consumers, the Joint FAO/WHO Expert Committee on Food Additives evaluated and provided the maximum residue limits (MRLs) for veterinary antibiotic residues permitted in food (5). The MRLs were determined based on acceptable daily intake (ADI) that should be harmless to humans according to results from extensive toxicity studies. However, these studies did not evaluate animal models of chronic metabolic diseases and were designed without considering the effects of antibiotic residues on gut microbiota and microbial metabolites, thereby failing to establish tolerable levels for the gut microbiota that can affect health of the host and cause disease.

Previous studies reported that antibiotics remained in meat, eggs, milk, and seafood products (6–8), sometimes even at levels exceeding the MRLs (9). Moreover, cooking processes, such as frying and roasting, can increase the concentrations of certain antibiotics (7), raising the probability of exposure to antibiotic residues through food. Therefore, veterinary antibiotics can be detected in human urine due to the consumption of pork, chicken, and dairy products (8). Furthermore, higher levels of veterinary antibiotics detected in the urine were reported to be positively correlated with obesity in children, revealing that exposure to antibiotic residues in food may contribute to obesity (10).

The commensal bacteria play a crucial role in human health and disease mainly by producing various metabolites, such as short-chain fatty acids (SCFAs) and secondary bile acids (11). Indeed, dysbiosis of the gut microbiota and microbial metabolites have been associated with metabolic diseases (12, 13). Antibiotics significantly disturb the composition of the gut microbiota, alter SCFAs and bile acids, as well as their signaling pathways, thereby leading to metabolic consequences (14–16). Hence, considering their importance in treating infections, antibiotics can be viewed as a double-edged sword for human health given their untoward effects on the gut microbiota and host metabolic homeostasis (17).

The gut microbial community is dynamic and susceptible to environmental shifts in early life, which has been considered the critical window of gut microbiota development (18). Clinical studies reported that antibiotic exposure during infancy is associated with increased risks of being overweight and obese (19–22). Moreover, studies in mice showed that early-life antibiotic exposure disturbs the colonization and maturation of the intestinal microbiota, leading to lasting effects on the metabolism of the host (23–25), even when the antibiotic is administrated at sub-therapeutic doses (18, 26, 27). The abovementioned evidence demonstrate that low-dose antibiotic exposure, especially in early life, is sufficient to induce undesirable metabolic consequences. Along with the findings of epidemiological studies, low dose veterinary antibiotic residues in food are believed to potentially promote obesity via gut microbiota perturbation. However, the impact of residual dose of antibiotics on the gut microbiota and human health has not been elucidated (3, 28).

In this study, the impact of ADI and theoretical maximum daily intake (TMDI) dosage of tylosin—used as a model antibiotic growth promoter owing to its frequent use and residual detection in food—on host metabolism and gut microbiota was explored in mice. ADI can be safely consumed daily over life without any appreciable health risk (29), whereas TMDI is the estimation of the maximal residual dose that can be consumed from foods according to MRL (5). To study the effect of tylosin-altered microbiota on obesity-related phenotypes, fecal samples were transplanted from mice fed TMDI dose of tylosin into germ-free mice. In addition, whether early-life exposure to TMDI dose of tylosin could induce obesity-related complications and cause alterations in gut microbiota and microbial metabolites was also investigated. Finally, a plausible mechanism by which tylosin TMDI dose induced metabolic consequences via altering bile acid composition and the ileal fibroblast growth factor 15 (FGF15)/hepatic fibroblast growth factor receptor 4 (FGFR4) pathway.

## Results

### Residual doses of tylosin facilitate obesity preferentially in high-fat diet-fed mice

To investigate whether chronic exposure to an acceptable or residual dose of antibiotic could cause obesity, and the potential synergistic effects of antibiotics and diet on host metabolism, a murine model that included two doses of antibiotic (tylosin at ADI and TMDI dose) and different diets (normal chow diet [NCD] and high-fat diet [HFD]) was designed **(Fig. 1a)**.

**FIG 1.**
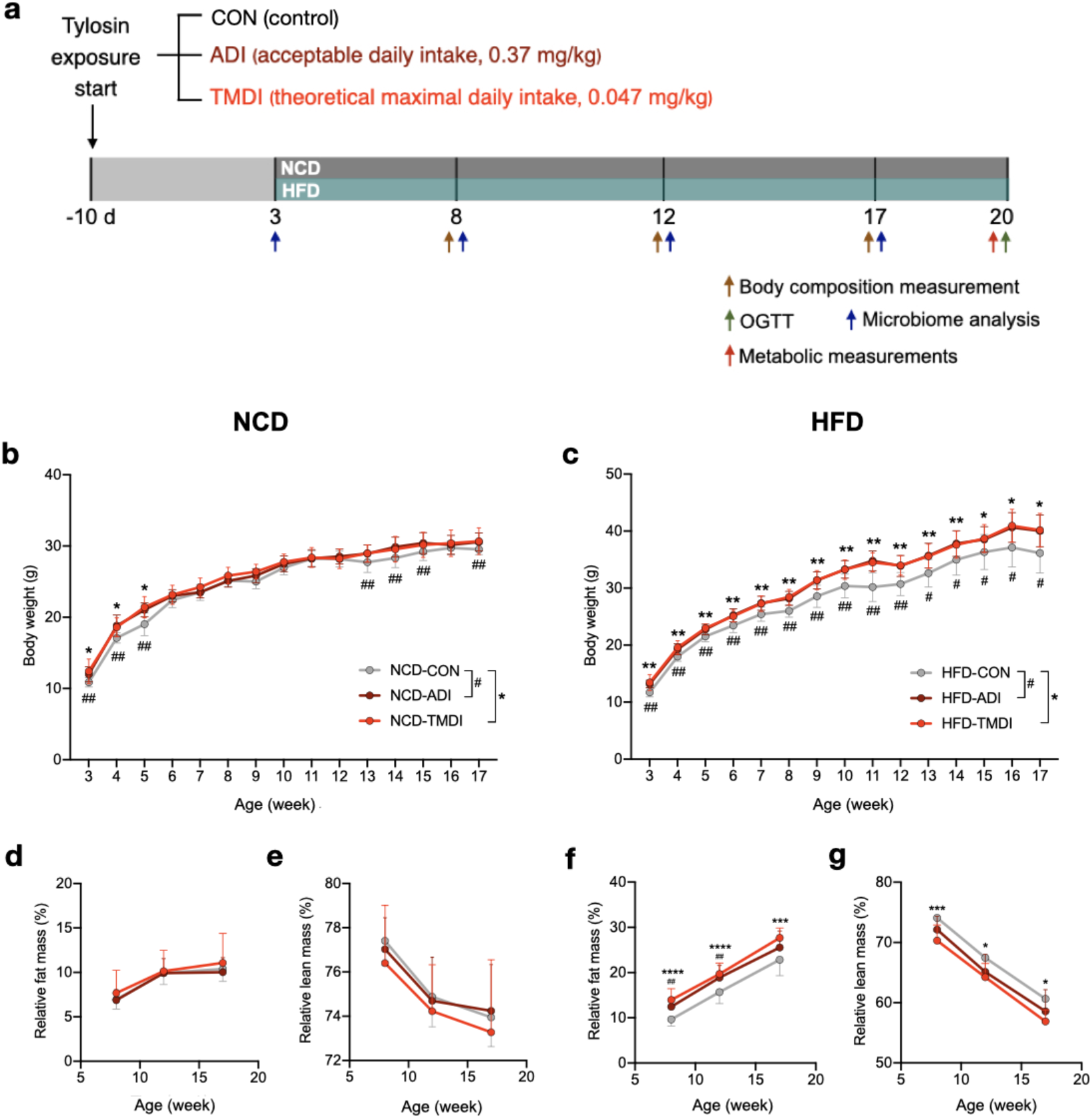
Residual dose of tylosin facilitates HFD-induced obesity. **(a)** Experimental design of antibiotic residue exposure model. Body weight, relative fat mass, and relative lean mass changes in **(b, d, e)** NCD and **(c, f, g)** HFD mice. Data are expressed as mean ± SD (n = 10–12). NCD-CON versus NCD-ADI and HFD-CON versus HFD-ADI (^#^*p* < 0.05; and ^##^*p* < 0.01) and NCD-CON versus NCD-TMDI and HFD-CON versus HFD-TMDI (* *p* < 0.05; ** *p* < 0.01; and *** *p* < 0.001) by one-way ANOVA with Tukey’s range test. Abbreviations: ADI, acceptable daily intake; AUC, area under curve; CON, control; HFD, high-fat diet; HOMA-IR, homeostatic model assessment of insulin resistance; NCD, normal chow diet; OGTT, oral glucose tolerance test; TMDI, theoretical maximum daily intake.

Compared with NCD-CON (non-exposed/control) mice, NCD-ADI and NCD-TMDI mice showed significantly greater weight gain from weaning to 5 weeks of age. NCD-ADI mice also showed more weight gain from weeks 13 to 15 **(Fig. 1b)**, but no significant changes in relative fat and lean mass were observed among the NCD groups **(Fig. 1d, e)**. In contrast, HFD-ADI and HFD-TMDI mice exhibited significantly increased weight gain compared with HFD-CON group throughout the experiment from weaning to 17 weeks of age **(Fig. 1c)**. Body composition analysis showed that HFD-ADI and HFD-TMDI mice had increased relative fat mass at weeks 8, 12, and 17 compared with HFD-CON mice **(Fig. 1f)**, while HFD-TMDI mice also had decreased relative lean mass **(Fig. 1g)**. These results demonstrate that tylosin-induced adiposity is evident early in life with both NCD and HFD, but continuous effect of tylosin-induced adiposity requires concomitant HFD.

### Residual doses of tylosin exacerbate HFD-induced hepatic steatosis, adiposity, and insulin resistance

Antibiotics have been shown to induce obesity, nonalcoholic fatty liver disease (NAFLD), and insulin resistance (30). Given that tylosin increased the fat mass in HFD-fed mice but not in NCD-fed mice, additional investigations in the HFD-fed mice were conducted.

HFD-TMDI mice showed increased visceral fat, including epididymal and perinephric adipose tissues **(Fig. 2a, b)**. HFD-ADI and HFD-TMDI mice also showed adipocyte hypertrophy (**Fig. 2c**) and elevated adipocyte size (**Fig. 2e**) compared with HFD-CON mice. Histological examination of the liver revealed that tylosin-treated mice exhibited increased lipid droplet formation (**Fig. 2d**), higher number of inflammatory foci, and fatty liver score **(Fig. 2f)**, suggesting that residue levels of tylosin caused more severe NAFLD. No change in plasma total triacylglycerol, cholesterol, and lipopolysaccharides (LPS) was observed **(Fig. S2b, c, and g)**. The oral glucose tolerance test (OGTT) indicated that HFD-ADI and HFD-TMDI mice exhibited a trend toward higher plasma glucose during OGTT **(Fig. S2a)**, OGTT_AUC_ **(Fig. 2g)**, and fasting glucose **(Fig. 2h)**. The fasting insulin and the homeostasis model assessment of insulin resistance (HOMA-IR) index were also elevated in HFD-ADI mice compared with HFD-CON mice **(Fig. 2i, j)**. These results reveal that even residues of tylosin can induce adverse effects on metabolism when an HFD is consumed.

**FIG 2.**
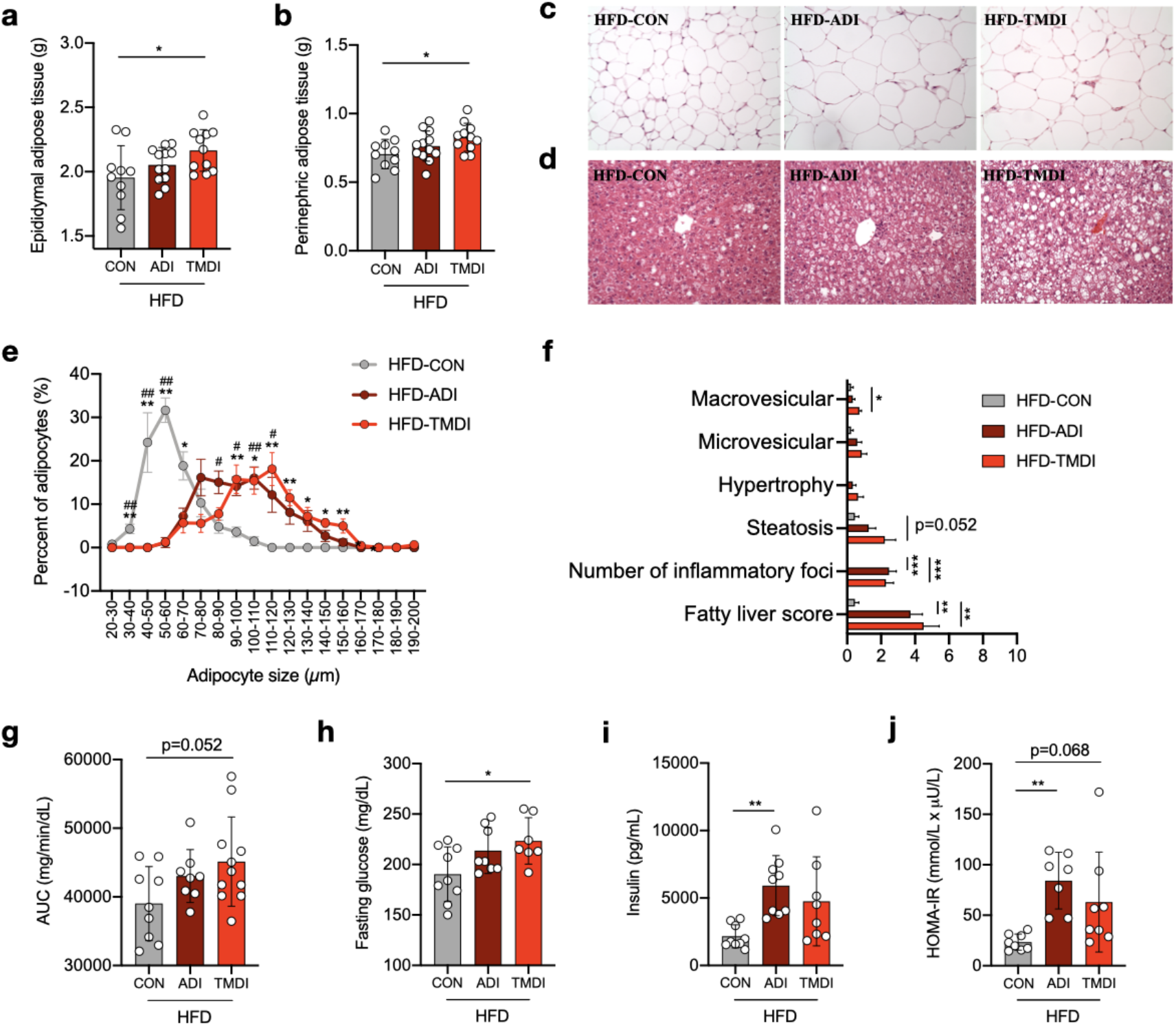
Residual dose of tylosin causes adiposity and insulin resistance. **(a)** Weight of epididymal adipose tissue. **(b)** Weight of perinephric adipose tissue. **(c-d)** histological features of H&E stained in **(c)** epididymal adipose tissue and **(d)** liver. **(e)** Adipocyte diameter of epididymal adipose tissue. **(f)** Fatty liver score including steatosis (macrovesicular, microvesicular, and hypertrophy) and inflammation (number of inflammatory foci). **(g)** Area under curve (AUC) derived from the OGTT. **(h)** Plasma glucose and **(i)** insulin level after overnight fasting. **(j)** HOMA-IR index represented as an indicator of insulin resistance. Data are expressed as mean ± SD (n = 8–12 mice per group). HFD-CON versus HFD-ADI (^#^*p* < 0.05 and ^##^*p* < 0.01) and HFD-CON versus HFD-TMDI (* *p* < 0.05; ** *p* < 0.01; and *** *p* < 0.001) by one-way ANOVA with Tukey’s range test. Abbreviations: ADI, acceptable daily intake; AUC, area under curve; CON, control; HFD, high-fat diet; HOMA-IR, homeostatic model assessment of insulin resistance; NCD, normal chow diet; TMDI, theoretical maximum daily intake.

### Early-life exposure to residual doses of tylosin alter the gut microbiota composition

Sub-therapeutic antibiotic treatment was previously shown to disrupt the development and maturation of the gut microbiota with metabolic consequences (18, 24, 25, 27, 30). Thus, sequencing of fecal *16S* rRNA was performed in mice at 3 (weaning), 8, and 17 weeks of age to investigate changes in the gut microbiota. Compared with HFD-TMDI and HFD-CON mice, HFD-ADI mice showed a significant reduction in the Shannon Index at 3 weeks that dramatically increased during weeks 3–17 **(Fig. 3a)**. Principal coordinate analysis (PCoA) based on Bray-Curtis dissimilarity revealed that the microbiomes of NCD and HFD mice were clustered separately **(Fig. S3a)**, indicating that diet may be the most important factor influencing the composition of the gut microbiota. Tylosin also influenced the gut microbiota composition in both NCD-fed and HFD-fed mice, with greater shifts when administrated at higher doses **(Fig. S3b, c)**. The effect of tylosin was most evident on the immature early-life gut microbiota at week 3, with gradual maturation and clustering of the gut microbiota at weeks 8 and 17, respectively (*p* < 0.05; **Fig. 3b, c and Fig. S3c–e)**. These results suggest that residual dose of tylosin impacts on the gut microbiota composition in early life, but the gut microbiota could subsequently mature and recover.

**FIG 3.**
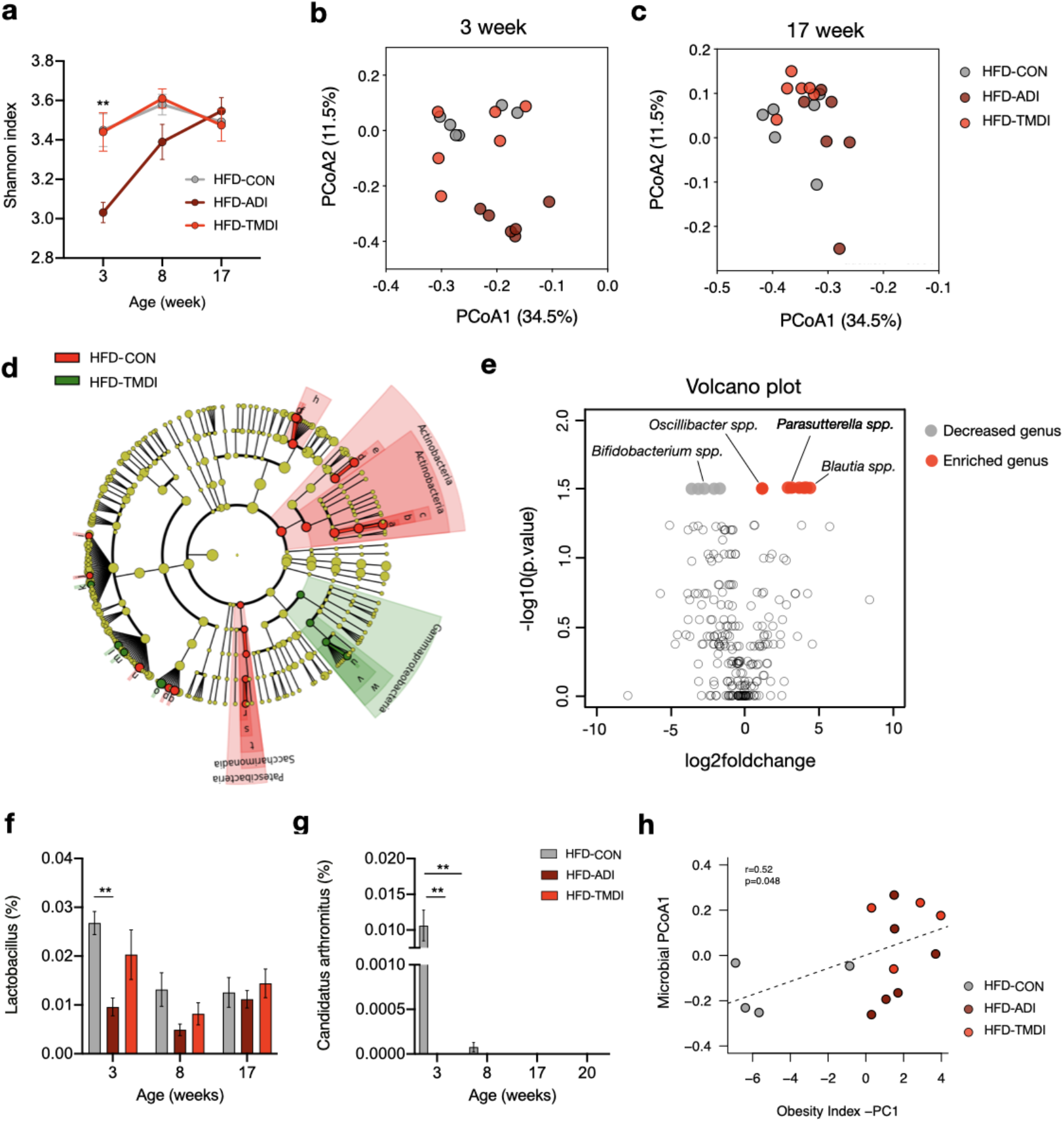
Residual dose of tylosin impacts on gut microbiota composition. **(a)** Change in Shannon index in 3, 8, 17 week-old mice. Principal coordinate analysis (PCoA) of Bray-Curtis distances of the microbiota at **(b)** 3 week-old and **(c)** 17 week-old. **(d)** Linear discriminant analysis effect size (LEfSe) showing enriched bacterial phyla. **(e)** Volcano plot showing enriched and decreased bacterial genera in HFD-TMDI group. Relative abundance of **(f)** *Lactobacillus* and **(g)** *Candidatus arthromitus*. **(h)** Spearman’s correlation of microbial PCoA1 index and the obesity index. Data are expressed as mean ± SEM (n = 8-12). * *p* < 0.05 and ** *p* < 0.01 by Wilcoxon signed-rank test. *p*-values of PCoA were assessed by ADONIS test. Abbreviations: ADI, acceptable daily intake; CON, control; HFD, high-fat diet; TMDI, theoretical maximum daily intake.

Linear discriminant analysis effect size (LEfSe) analysis and volcano plot were performed to elucidate the relative abundances of bacterial taxa that significantly differed with and without tylosin exposure. *Gammaproteobacteria*, comprising bacterial taxa generally regarded as LPS-producing pathogens (31), were more abundant in both HFD-ADI and HFD-TMDI groups than in HFD-CON group **(Fig. 3d and Fig. S4a)**. *Tyzzerella*, reported as a key bacterial taxon associated with infant antibiotic exposure-related obesity, was enriched in HFD-TMDI mice (20). In turn, bacterial taxa within the beneficial *Actinobacteria* phylum, particularly *Bifidobacterium* species, were more abundant in HFD-CON mice **(Fig. S4b)**. Volcano plot revealed that *Oscillibacter* spp., which has been associated with obesity and intestinal permeability (32), was significantly enriched in HFD-TMDI mice, whereas *Bifidobacterium* spp. was significantly reduced in HFD-TMDI mice **(Fig. 3e)**. Moreover, *Lactobacillus* and *Candidatus arthromitus*, which have been associated with obesity prevention (18), were significantly decreased in HFD-ADI and HFD-TMDI mice **(Fig. 3f, g)**. These results demonstrate that TMDI dose of tylosin depleted the beneficial bacteria and enriched the pathogenic bacteria. Correlation analysis between the beta diversity (PCoA1 index) of gut microbiota and obesity index (PC1 based on the obesity-related parameters; **Fig. 3h**) exhibited a positive correlation, suggesting that the gut microbiota alteration is involved in the obesity and metabolic disorder phenotypes.

### Fecal microbiota transplantation from HFD-TMDI mice induce adiposity and insulin resistance in germ-free mice

To further investigate whether residual tylosin exposure could facilitate obesity through the alternation of gut microbiota, fecal microbiota transplantation (FMT) was performed **(Fig. 4a)**. Considering that TMDI dose can better simulate human exposure to antibiotics in food, feces from HFD-TMDI mice were transplanted to germ-free mice.

**FIG 4.**
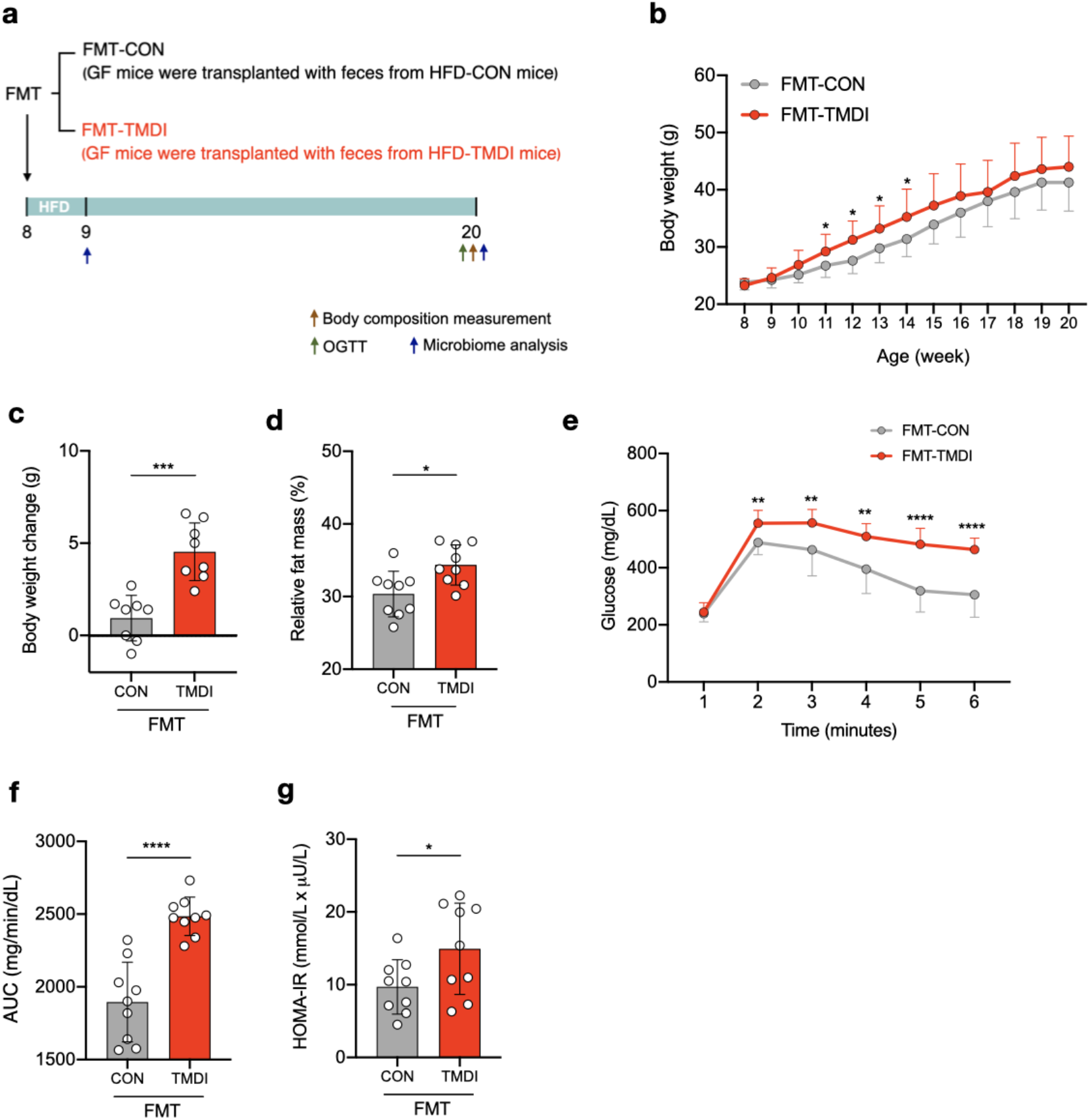
Fecal microbiota transplantation of HFD-TMDI feces induces obesity and insulin resistance in germ-free mice. **(a)** Experimental design of the FMT study. **(b)** Body weight change. **(c)** Body weight change at 2 weeks after FMT (from 8–10 weeks of age). **(d)** Body fat mass at 20 weeks of age. **(e)** Plasma glucose profile measured during the OGTT. **(f)** AUC derived from the OGTT. **(g)** HOMA-IR index. Data are expressed as mean ± SD (n = 8–10). * *p* < 0.05; ** *p* < 0.01; *** *p* < 0.001; and **** *p* < 0.0001 by un-paired *t*-test. Abbreviations: AUC, area under curve; CON, control; FMT, fecal microbiota transplantation; HOMA-IR, homeostatic model assessment of insulin resistance; OGTT, oral glucose tolerance test; TMDI, theoretical maximum daily intake.

Compared with germ-free mice that received feces from HFD-CON mice (FMT-CON), the HFD-TMDI recipient mice (FMT-TMDI) showed higher body weight at 11, 12, 13, 14 weeks of age **(Fig. 4b)**, increased weight gain during weeks 8–10 **(Fig. 4c)**, and elevated fat mass at week 20 **(Fig. 4d)**, indicating that the microbiome from HFD-TMDI mice increased the adiposity of the recipient mice. Additionally, HFD-TMDI mice exhibited increased plasma glucose during OGTT, OGTT_AUC_, and HOMA-IR index **(Fig. 4e-g)**. The intestinal permeability measurement by fluorescein isothiocyanate-dextran revealed increased permeability in FMT-TMDI mice, although differences in plasma LPS levels were not observed between the two groups. **(Fig. S5f, g)**. Thus, TMDI dose of tylosin-altered microbiota induced the metabolic phenotype of obesity and insulin resistance in germ-free recipient, indicating that the microbiota play a causative role in tylosin TMDI-induced metabolic disorders.

### Exposure to TMDI dose of tylosin in early life is sufficient to cause lasting metabolic complications

Accumulating evidence indicates that antibiotic exposure in early life, which is the critical developmental period of gut microbiota, causes obesity later on (19–22). Results of our first experiment showed that 3-week-old HFD-TMDI mice exhibited increased body weight **(Fig. 1b, c)**, indicating that early-life exposure to residual dose of tylosin induces weight gain. Accordingly, an early-exposure experiment was conducted to investigate the influence of early-life exposure to tylosin TMDI on obesity-related phenotypes and the gut microbiota **(Fig. 5a)**.

**FIG 5.**
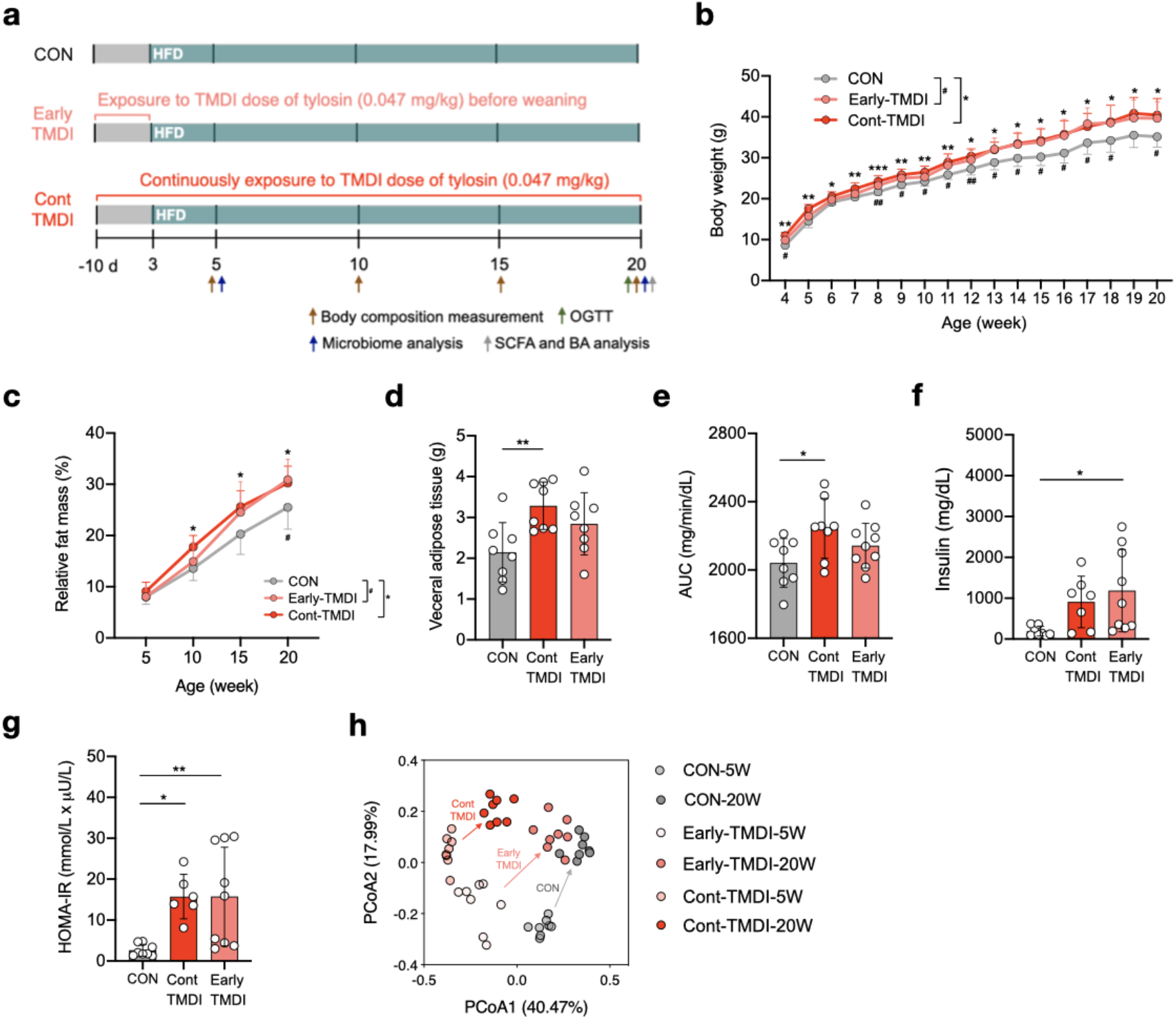
Early-life exposure of TMDI dose of tylosin induces metabolic disorders. **(a)** Experimental design of the early life exposure model. **(b)** Body weight change. **(c)** Relative fat mass. **(d)** Weight of visceral adipose tissue. **(e)** AUC derived from the OGTT. **(f)** Insulin level after overnight fasting. **(g)** HOMA-IR index. **(h)** PCoA of Bray-Curtis distances of the microbiota at 5 and 20 weeks of age. Data are expressed as mean ± SD (n = 6–10). Statistical analysis of **(b)** and **(c)** were performed by one-way ANOVA with Tukey’s range test comparing CON versus Early-TMDI (^#^p < 0.05 and ^##^*p* < 0.01) or CON versus Cont-TMDI (* *p* < 0.05; ** *p* < 0.01; and *** *p* < 0.001) at each time point. Statistical analyses of **(d–g)** were performed by one-way ANOVA with Tukey’s range test (* *p* < 0.05 and ** *p* < 0.01). *p*-values of PCoA **(h)** were assessed by ADONIS test. Abbreviations: AUC, area under curve; BA, bile acid; CON, control; HFD, high-fat diet; HOMA-IR, homeostatic model assessment of insulin resistance; OGTT, oral glucose tolerance test; SCFA, short-chain fatty acid; TMDI, theoretical maximum daily intake. Treatment regimen: Cont-TMDI, continuous exposure to TMDI dose of tylosin; Early-TMDI, exposure to TMDI dose of tylosin early in life.

Cont-TMDI mice, which were continuously exposed to tylosin TMDI throughout the experimental period, displayed continuously elevated body weight, relative fat mass **(Fig. 5b, c)**, weight of visceral fat mass, and OGTT_AUC_ **(Fig. 5d, e)** compared with HFD-CON. Remarkably, Early-TMDI mice, which were exposed to tylosin TMDI during pregnancy and nursing period, also showed a constant increase in body weight, relative fat mass, fasting insulin, and HOMA-IR **(Fig. 5b-g)** after cessation of tylosin exposure. These findings suggest that exposure to residual dose of antibiotics from food early in life, can have long-lasting effect on metabolism and lead to obesity.

### Early exposure to TMDI dose of tylosin alter the abundance of specific bacteria related to the metabolic homeostasis of the host

The overall gut microbiota composition based on PCoA of Bray-Curtis distances showed that tylosin influenced the gut microbiota composition at weeks 5 and 20 (*p* < 0.05) **(Fig. 5h)**. Compared with CON mice, Cont-TMDI mice showed a greater difference than Early-TMDI mice, suggesting that continuous exposure to tylosin modified the overall gut microbiota composition more than exposure restricted to early life. Furthermore, Early-TMDI vs. CON group exhibited more differences at week 5 than at week 20, possibly due to maturation of the gut microbiota **(Fig. 5h)**.

Since both Early-TMDI and Cont-TMDI mice displayed shifts in microbial composition at 20 weeks of age (*p* < 0.05), bacterial taxa that differed in relative abundances from CON mice were identified and examined their association with obesity-related phenotypes. Overall, 32 bacterial genera were found to be significantly altered by early or continuous exposure to tylosin TMDI **(Fig. 6)**. Despite discontinuation of tylosin exposure in Early-TMDI mice at 3 weeks of age, several bacterial genera remained altered at week 20. Interestingly, bacterial genera that were increased in both Early-TMDI and Cont-TMDI mice, including *Anaerofustis*, demonstrated a significant positive correlation with obesity-related phenotypes **(Fig. 6)**. In contrast, genera that were depleted in both Early-TMDI and Cont-TMDI, including bacteria within *Lachnospiraceae* and *Ruminococcaceae* families, exhibited a significant negative correlation with obesity-related phenotypes **(Fig. 6)**. These findings indicate that exposure to residual dose of tylosin influences the abundance of specific bacteria involved in the regulation of the metabolic homeostasis of the host, even when the exposure is limited to early life.

**FIG 6.**
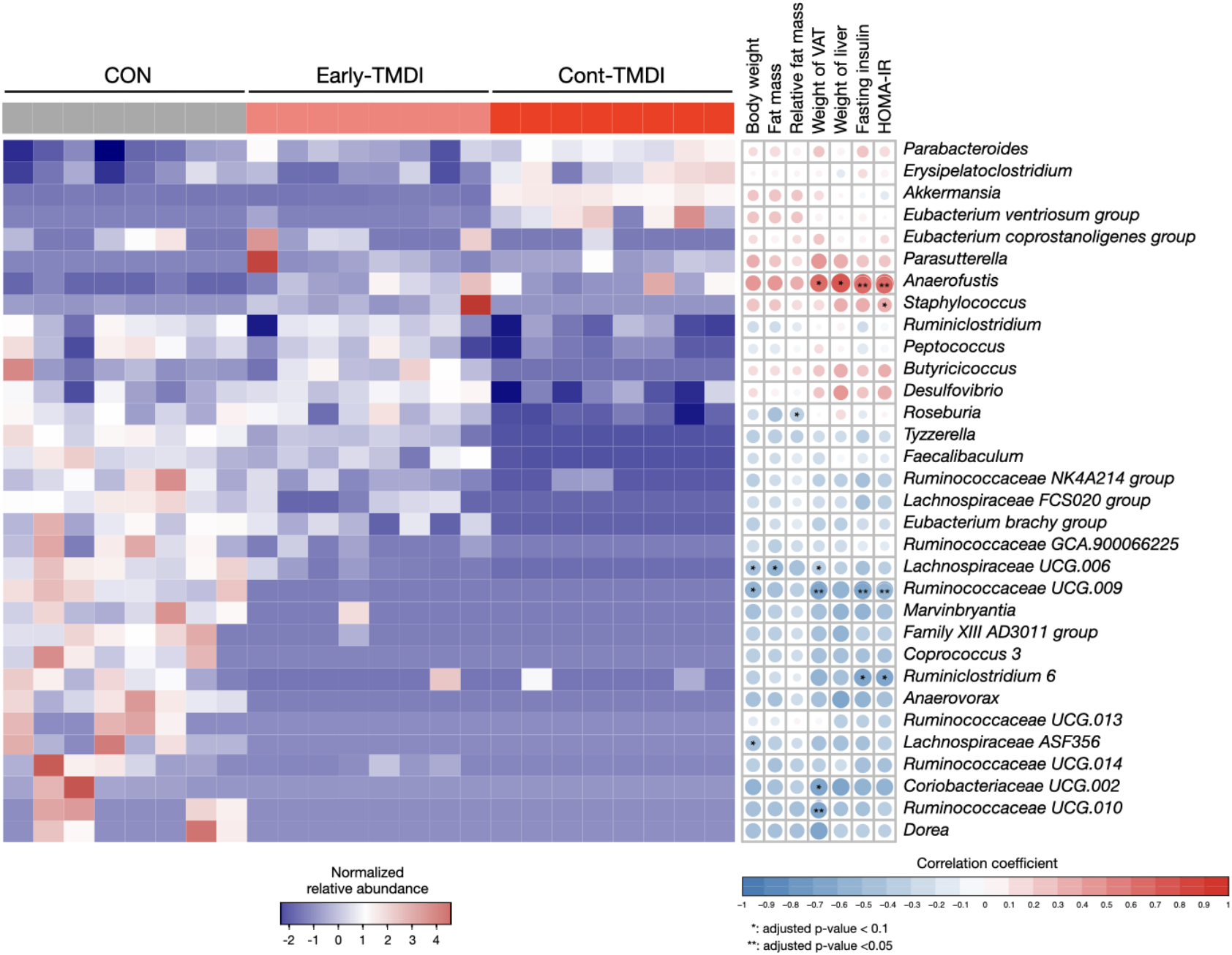
TMDI of tylosin alters the abundance of specific bacteria with correlation to host metabolic phenotype. **(Left panel)** Heatmap of significantly enriched or reduced bacterial genera in CON, Early-TMDI, and Cont-TMDI groups, with genus name shown to the right of each row. Log10-transformed relative abundance is scaled and color-coded with index shown beneath the heatmap. Statistical analyses were performed by Kruskal—Wallis test (*q* < 0.05). **(Right panel)** Obesity and insulin resistance-related phenotypes correlated with bacterial genera. The size and color of the symbols represent the Spearman’s correlation coefficients. Abbreviations: CON, control; TMDI, theoretical maximum daily intake; VAT, visceral adipose tissue. Treatment regimen: Cont-TMDI, continuous exposure to TMDI dose of tylosin; Early-TMDI, exposure to TMDI dose of tylosin early in life.

### Exposure to TMDI dose of tylosin alter the composition of short-chain fatty acids and the conversion of bile acids with downstream effects on the FGF15 signaling pathway

We further investigated the effect of TMDI dose of tylosin on major microbial metabolites including SCFAs and bile acids, which play a role in metabolic homeostasis by affecting multiple receptors and downstream signaling pathways. Propionic acid and butyric acid, two main SCFAs, were reduced in Cont-TMDI mice **(Fig. 7a)**. The duration of tylosin exposure influenced SCFA composition. For example, Early-TMDI mice only exhibited significant reduction in isobutyrate, whereas Cont-TMDI mice exhibited significant reductions in both branched-chain isobutyric acid and isovaleric acid **(Fig. 7a)**.

**FIG 7.**
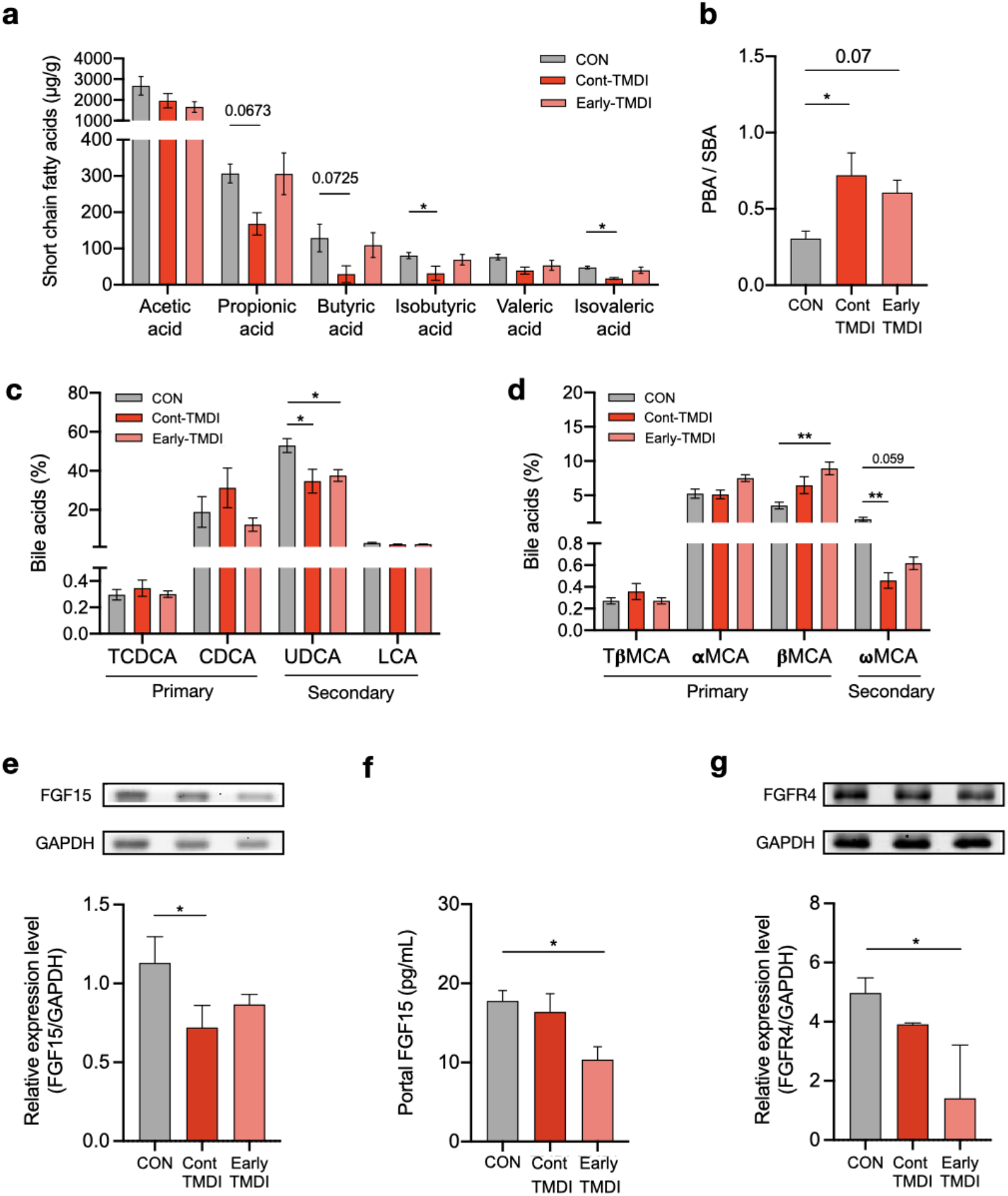
TMDI dose of tylosin decreases SCFA levels, increases PBA/SBA ratio, and alters downstream expressions of FGF15 and FGFR4. **(a)** Fecal SCFA levels. **(b)** Ratio of PBA to SBA. **(c)** Levels of non-12-OH bile acids. **(d)** Levels of 12-OH bile acids. **(e)** Western blotting of ileal FGF15 expression normalized to GAPDH. **(f)** FGF15 level in portal vein measured by enzyme-linked immunosorbent assay. **(g)** Western blotting of hepatic FGFR4 level normalized to GAPDH. Data are presented by mean ± SEM (n = 8–10) in **(a-d)** and mean ± SD (n = 4) in **(e-g)**. Statistical analyses were performed by one-way ANOVA with Tukey’s range test (* *p* < 0.05 and ** *p* < 0.01). Abbreviations: α-MCA, α-muricholic acid; β-MCA, β-muricholic acid; ω-MCA, ω-muricholic acid; CON, control; CDCA, chnodeoxycholic acid; FGF15, fibroblast growth factor 15; FGFR4, fibroblast growth factor receptor 4; GAPDH, glyceraldehyde 3-phosphate dehydrogenase; LCA, lithocholic acid; PBA, primary bile acid; SBA, secondary bile acid; T-β-MCA, tauro-beta-muricholic acid; TCDCA, taurochenodeoxycholic acid; TMDI, theoretical maximum daily intake; UDCA, ursodeoxycholic acid. Treatment regimen: Cont-TMDI, continuous exposure to TMDI dose of tylosin; Early-TMDI, exposure to TMDI dose of tylosin early in life.

The total amount of bile acids and the ratio of conjugated to unconjugated bile acids were not changed by tylosin **(Fig. S7a and b)**. However, the ratio of primary bile acids (PBA) to secondary bile acids (SBA) was significantly increased in Cont-TMDI mice, with a trend toward increase in Early-TMDI mice **(Fig. 7b)**. A correlation analysis between the PBA/SBA ratio and obesity index showed a moderate trend **(Fig. S7h)**. Detected bile acids were classified based on their metabolic pathway (33) into non-12-OH bile acids **(Fig. 7c)**, muricholic acids (MCA; **Fig. 7d)**, and 12-OH bile acids **(Fig. S7c**). *β*-MCA **(Fig. 7d)**, the unconjugated PBA, was significantly increased, whereas ursodeoxycholic acid and ω-MCA **(Fig. 7c, d)**, the unconjugated SBA, were decreased in tylosin-treated mice, possibly due to the inhibition of bacteria involved in epimerization and dihydroxylation of PBAs, such as *Clostridia, Peptostreptococcus, Bifidobacterium,* and *Lactobacillus* **(Fig. S7d–g)** (34). These results revealed that the increased PBA/SBA ratio caused by tylosin TMDI might be related to obesity.

Given that the alternation of PBA/SBAa ratio contributes to plasma FGF15, an insulin-like hormone secreted in the ileum (35, 36), and regulates hepatic lipid and glucose metabolism by binding to FGFR4 (37, 38), the FGF15/FGFR4 signaling pathway was explored next. Tylosin was found to inhibit the conversion of PBA to SBA, and reduce ileal FGF15 expression **(Fig. 7e and fig. S7i)** and portal vein FGF15 levels **(Fig. 7f)** with subsequent reduction in hepatic FGFR4 **(Fig. 7g)**; thus, potentially affecting hepatic insulin sensitivity and lipid metabolism. Collectively, these results showed that TMDI dose of tylosin alter the intestinal microbiome and bile acid metabolism with downstream effects on the farnesoid X receptor signaling pathway, decreasing bile acid-related FGF15 levels and leading to obesity-related metabolic dysfunction **(Fig. 8)**.

**FIG 8.**
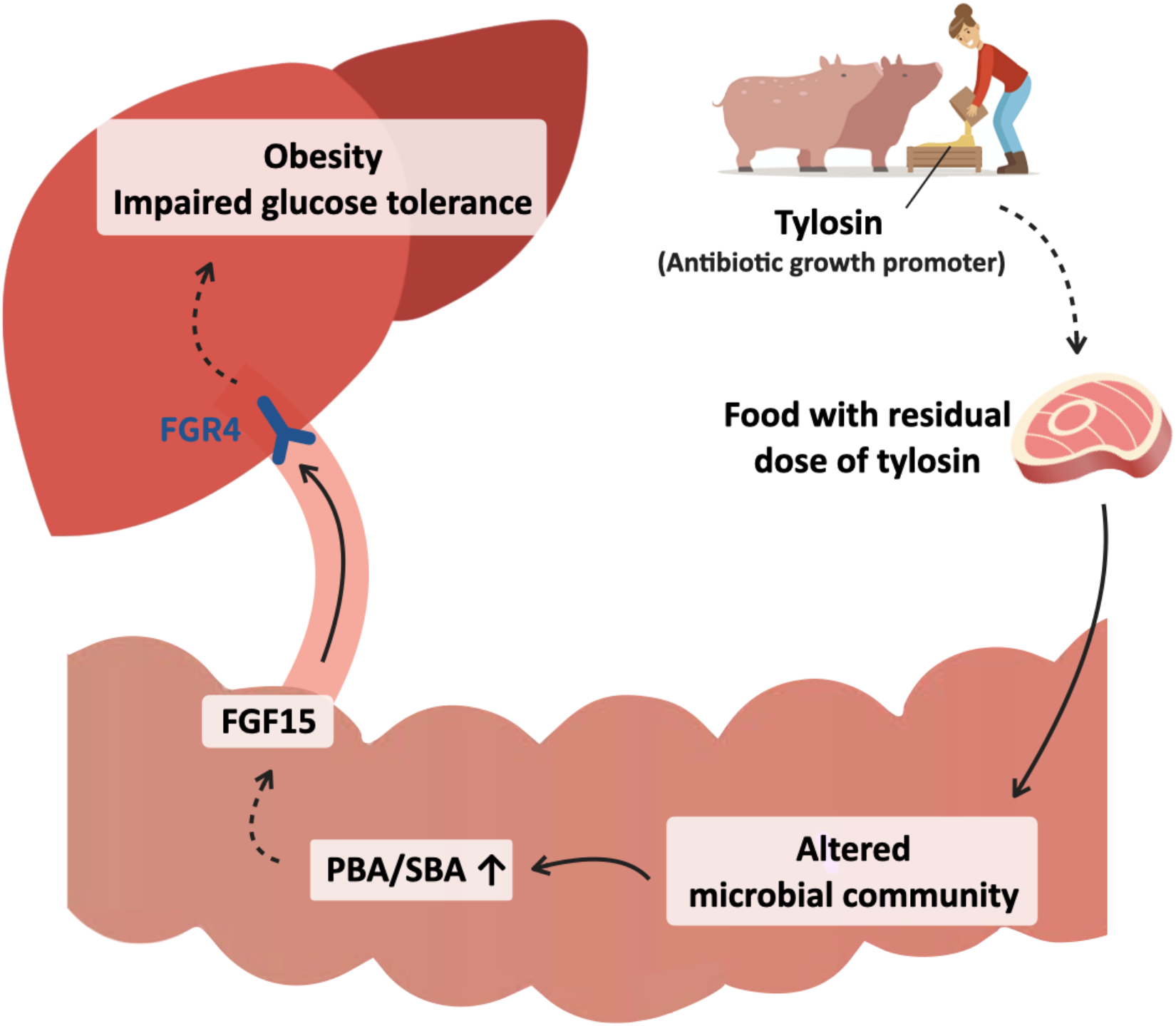
Residual dose of antibiotic growth promoter exacerbates HFD-induced metabolic disorder by altering the gut microbiota, microbial metabolites, and downstream signaling pathway. Abbreviations: AGP, antibiotic growth promoter; FGF15, fibroblast growth factor 15; HFD; high fat diet; PBA, primary bile acid; SBA, secondary bile acid.

## Discussion

Previous studies indicated that sub-therapeutic antibiotic treatment and low-dose penicillin led to obesity and NAFLD (18, 27, 30). The TMDI dose, which is used for simulating the ingestion of antibiotic residues through food consumption in the present study, was 20-fold lower than low-dose penicillin and 425-fold lower than the growth-promoting dose used in food animals (18, 39). Herein, ADI and TMDI doses of tylosin were shown to induce obesity, fatty liver, and insulin resistance in HFD mice **(Figs. 1 and 2)**. Despite its extremely low dose, tylosin TMDI significantly enhanced fat accumulation and insulin resistance, suggesting that continuous exposure to very low dose antibiotic residues in food can affect human metabolism. Interestingly, mice administered tylosin on the NCD did not exhibit increased fat mass. One plausible explanation is that tylosin amplified the effect of dysbiosis caused by the HFD. Moreover, Early-TMDI mice exhibited increased weight, fat mass, and insulin resistance index at 20-week old, despite the administration of TMDI dose of tylosin was ceased at weaning. Hence, these findings suggest that early exposure to antibiotic residue has a long-lasting effect on metabolism.

The diversity analysis of gut microbiota observed indicates a dose-dependent effect of tylosin on the gut microbiota composition **(Fig. 3a–c)** and reveals that continuous tylosin exposure has a more significant effect compared to exposure restricted to early life **(Fig. 5h)**. Interestingly, compared with CON mice, Early-TMDI mice exhibited a more similar microbial community at week 20 than at week 5 based on PCoA **(Fig. 5h)**, but fat mass was more significantly increased at 20 weeks of age **(Fig. 5c)**. This finding suggests that the impaired metabolic phenotype can persist despite recovery of the microbiota, consistent with findings from previous investigations (18). In addition, minor disruption of the microbiota seems to be sufficient for inducing significant adiposity (24).

The altered abundance of specific bacterial taxa correlates with obesity-related phenotypes **(Fig. 6)**. Overall, genera enriched in Early-TMDI and Cont-TMDI mice were found to be positively correlated with obesity and insulin resistance, whereas those depleted in Early-TMDI and Cont-TMDI mice were negatively correlated. These bacteria have been found to be related to obesity in previous studies. For instance, *Anaerofustis*, the tylosin-enriched bacterium, was found to be increased in obese humans (40). In turn, the tylosin-depleted *Ruminococcaceae* and *Lachnospiraceae* were found to be associated with a lower longitudinal weight gain (41). These findings indicate that despite discontinuation of TMDI-tylosin at 3 weeks of age, several genera related to host metabolism remained altered at 20 weeks of age. Therefore, colonization could be perturbed by exposure to a very low dose of antibiotic residue early in life, which can in turn contribute to metabolic disorders in adulthood (18, 26, 42, 43).

Herein, a metabolomic analysis revealed that TMDI dose of tylosin modified both SCFAs and bile acids. Cont-TMDI mice showed decreased isobutyric acid and isovaleric acid, which were reported to improve insulin-stimulated glucose uptake and enhance insulin sensitivity (44). Additionally, Cont-TMDI mice showed a reducing trend of butyric acid and propionic acid, which were associated with enhanced intestinal barrier function and insulin sensitivity (11, 45). Reduction of SCFAs could have contributed to the depletion of *Lachnospiraceae* and *Ruminococcaceae* in Early-TMDI and Cont-TMDI mice (40), and the observed increased PBA/SBA ratio may be caused by the broad inhibition of specific bacteria related to the conversion of PBA **(Fig. 6)**.

Increased PBA/SBA ratio has been associated with decreased insulin sensitivity in patients with NAFLD or nonalcoholic steatohepatitis (46, 47). In this study, tylosin TMDI-treated mice showed increased PBA/SBA ratio, lower FGF15 levels in the ileum and portal vein, thereby decreasing the expression of hepatic FGFR4, which may cause metabolic disorders by affecting metabolism-related signaling pathways in the liver (37, 38, 48). Consistent with these observations, antibiotics were reported to increase the PBA/SBA ratio, with subsequently decreased plasma FGF19 (human orthologue of FGF15) and peripheral insulin sensitivity (14).

This study has some limitations. The *in vivo* effects of tylosin, a macrolide antibiotic growth promoter, on the gut microbiota and the obesity phenotype were demonstrated. However, exposure to alternative antibiotics may lead to different outcomes owing to the specific antimicrobial action and spectrum of each antibiotic. Moreover, the human dietary pattern is dynamic, exposing us to multiple types of antibiotics, and even pesticides, in our daily lives. Future research should investigate the effects of other antibiotics at residual amounts and combination of different antibiotics to better reflect real-life conditions.

In conclusion, tylosin at ADI and TMDI doses, which are generally regarded as harmless, was shown to promote increased body weight, fat mass, and insulin resistance index in HFD-fed mice, and alter the gut microbiota composition. Moreover, altered gut microbiota was found to be critical for tylosin TMDI-induced metabolic consequences. Early-life exposure to TMDI dose of tylosin is sufficient to induce metabolic disorders, alter the abundance of specific bacteria related to host metabolic homeostasis, and modify the SCFA and bile acid composition. Lastly, exposure to TMDI dose of tylosin, whether continuously or restricted in early life, was shown to support lasting metabolic consequences via the ileal FGF15/hepatic FGFR4 pathway. Taken together, these findings indicate that the permissible exposure level of antibiotic residue should be re-established while considering its impact on the gut microbiota, for which this study provides valuable clues.

## Materials and Methods

### Antibiotic selection and dose calculation

Tylosin was selected as a model antibiotic growth promoter because of its high consumption levels in the annual consumption of veterinary antibiotics (Bureau of Animal and Plant Health Inspection and Quarantine, 2014). The doses of ADI and TMDI were obtained from the World Health Organization Technical Report Series (49). The ADI dose is derived from no observed adverse effect level, which is the lowest concentration that generates an adverse effect in long-term toxicity studies. The TMDI is an estimate of dietary intake obtained by MRLs and the sum of average daily per capita consumption of each food commodity (5).

### Experimental design of the animal studies

Animal experiments were performed with permission from the Institutional Animal Care and Use Committee of National Taiwan University (approval number: NTU-106-EL-051and NALC 107-0-006-R2). All mice were purchased from the National Laboratory Animal Center (Taipei City, Taiwan).

#### I. Antibiotic residue exposure model (Fig. 1a)

Pregnant mice were administered tylosin at the doses of ADI (0.37 mg/kg) and TMDI (0.047 mg/kg) through drinking water from day 10 of gestation. Control mice (CON) did not receive antibiotics. After the offspring were weaned, they were randomly divided into normal chow diet (NCD) (MFG, Oriental Yeast, Japan) and high-fat diet (HFD, 60% kcal from fat) (D12492, Research Diets, New Brunswick, NJ, USA). Body composition was measured at 8, 12, and 17 weeks. The metabolic measurements and OGTT were performed before euthanizing the mice at 20 weeks. After euthanasia the mice, the blood and tissue samples were collected and stored at −80 °C.

#### II. Fecal microbial transplantation study (Fig. 4a)

Eight-week-old C57BL/6 germ-free mice were randomly divided into fecal microbiota transplantation-control (FMT-CON) and FMT-TMDI groups, which were transplanted with the fecal microbiota from HFD-CON and HFD-TMDI mice, respectively. After FMT, the recipient mice were housed in two independent isolators and fed with irradiated HFD until 20 weeks of age. Before the recipient mice were euthanized, the body composition analysis and OGTT were performed.

#### III. Early-life exposure model (Fig. 5a)

The mice were divided into three groups: CON, feeding condition was the same as HFD-CON; Early-TMDI, exposure duration was limited during gestation and lactation period; Cont-TMDI, feeding condition was the same as HFD-TMDI. The experiment started on day 10 of gestation. Before the mice were weaned, both Early-TMDI and Cont-TMDI mice were exposed to TMDI dose of tylosin. After weaning, only the Cont-TMDI mice were continuously exposed to tylosin through the drinking water. Body composition was measured at 5, 10, 15, and 20 weeks. The OGTT was performed before euthanizing the mice at 20 weeks.

### Body composition analysis

Body composition was determined using Minispec LF50 TD-NMR Body Composition Analyzer (Bruker, Billerica, MA, USA), which provides the measurement of body weight, and lean and fat mass. The relative fat mass was calculated as fat mass (g)/body weight (g) ratio.

### Glucose and insulin sensitivity

For the OGTT, mice were deprived of food for 5 h. Blood was collected from the submandibular vein, and the glucose levels were measured by the glucometer (Roche, Basel, Switzerland) at 0, 15, 30, 60, 90, and 120 min after oral administration with 2 g/kg glucose. The fasting insulin levels were detected by an enzyme-linked immunosorbent assay (ELISA) kit (Mercodia, Uppsala, Sweden). The HOMA-IR index was calculated using the formula: fasting glucose (nmol/L) × fasting insulin (μU/mL)/22.5 (50).

### Histopathological analysis of liver and adipose tissue

Liver and adipose tissue sections were dissected and fixed in 10% formalin solution. Histopathological analysis was performed by the formalin-fixed, paraffin-embedded, and hematoxylin and eosin (H&E)-stained slide. The fatty liver score was estimated by a pathologist as previously described (51). The fatty liver score included the evaluation of steatosis (macrovesicular, microvesicular, and hypertrophy) and inflammation (number of inflammatory foci). The visceral adipocyte quantification was performed by HCImage Live software (HCImage, Sewickley, PA, USA).

### Obesity index

The obesity index was extracted from multidimensional phenotype measurements of mice, including total fat mass (g), body fat (%), average growth rate, weights of adipose tissues and liver, fatty liver score, fasting glucose, fasting insulin, HOMA-IR index, fasting triglycerides and fasting total cholesterol based on a principal component analysis algorithm (30).

### Statistical analysis

Data were represented as mean ± standard deviation (SD) or mean ± standard error of the mean (SEM). One-way analysis of variance (ANOVA) with Tukey’s range test or Student’s *t*-test was applied for intergroup comparisons. Statistical assessment of the gut microbiome, SCFAs, and bile acids was performed using the Kruskal–Wallis with/without false discovery rate or one-way ANOVA with Tukey’s range test. All statistical data were analyzed using Prism software (version 8.4.3; GraphPad Software, San Diego, CA, USA) or RStudio (version 1.2.5001, RStudio, Boston, MA, USA).

## Acknowledgements

This study was funded by the Ministry of Science and Technology (MOST), Taiwan (MOST 106-3114-B-002-003, MOST 107-2321-B-002-039, MOST 108-2321-B-002-051, MOST 109-2327-B-002-005, and MOST 109-2314-B-002-064-MY3). We would like to acknowledge the technical support provided by the following institutes: Fecal microbiota transplantation, isolator service and metabolic measurement were conducted by National Laboratory Animal Center, National Applied Research Laboratories, Taiwan. Analysis of bile acids was supported by the National Taiwan University Consortia of Key Technologies. Analysis of short chain fatty acid was supported by the ‘Center of Precision Medicine’ from The Featured Areas Research Center Program by the MOST. Body composition analysis was performed by Taiwan Mouse Clinic, Academia Sinica and Taiwan Animal Consortium.

## Author contributions

R.A.C. designed and performed the animal study, performed metabolomic, bioinformatics, and statistical analysis, and drafted the manuscript; W.K.W. proposed, designed, and instructed the study; S.P. assisted the experiments and bioinformatics analysis; P.Y.L. instructed the bioinformatics analysis; H.L.C. performed the germ-free mice study and assisted the metabolic measurement; Y.H.C. assisted the germ-free animal study and performed the LPS analysis; Q.L. instructed the bile acid analysis; H.C.H. and H.B.Z. performed SCFA analysis and assisted with mass spectrometry analysis; T.L.L. instructed the LPS analysis; Y.T.Y. conducted the PCR and library preparation for *16S* rRNA sequencing; H.S.H. and Y.E.L. instructed the western blotting; S.P., T.C.D.S., W.K.W., P.Y.L. and L.Y.S. critically revised the manuscript; W.K.W., L.Y.S., C.C.H., M.S.W., H.C.L., C.C.C. and C.T.H., provided professional insights, techniques, and relevant resources for the study.

## Data availability

The raw *16s* rRNA sequencing data are accessible at the National Center for Biotechnology Information Short Read Archive (BioProject: PRJNA715326).

