## Supplementary methods for "Dietary Exposure to Antibiotic Residues Facilitates Metabolic Disorder by Altering the Gut Microbiota and Bile Acid Composition"

***Corresponding authors:**

Plasma samples were centrifuged and added to HEK-Blue™ mTLR4 cell lines in HEK-Blue™ Detection medium (InvivoGen, CA, USA). After 24 h of incubation, the color of secreted embryonic alkaline phosphatase (SEAP) released by the reporter cells was measured by a spectrophotometer at 620 nm (1).

***Metabolic measurements***

Metabolic measurements (food intake, locomotor activity, VO2 consumption and VCO2 production) were obtained using the Promethion metabolic phenotyping system (Sable Systems, Las Vegas, NV, USA). Monitoring was performed for 24–48 h with ad libitum access to food and water after mice acclimatized to cages for 6–12 h.

***Fecal microbiota extraction and 16S rRNA gene sequencing***

Mice feces were collected and were frozen immediately at −80 ℃. Fecal microbial DNA was isolated by using QIAamp 96 PowerFecal QIAcube HT Kit (QIAGEN, MD, USA). The 16S V4 region was amplified by the forward primer (F515, 5´-GTGCCAGCMG CCGCGGTAA-3´) and the reverse primer (R806, 5´-GGACTACHVGGGTWTCTAAT-3´). The reaction conditions were 3 min at 95 °C, followed by 25 cycles of 95 °C for 30 s, 55 °C for 30 s, 72 °C for 30 s, and 5 min at 72

°C for a final extension. For library construction, the amplified PCR product was attached with Illumina sequencing adapters by Nextera XT Index Kit then purified by AMPure XP beads. The library quantification was conducted by Bioanalyzer DNA 1000 chip and Agilent Technologies 2100 Bioanalyzer. The sequencing (paired-end reads, 2 × 150 bp) was performed by Illumina NextSeq (Illumina, CA, USA).

***Bioinformatic analysis for microbial taxonomic profiling***

The amplicon sequences were processed by QIIME 2 pipeline (version 2019.10). The primer sequences of raw reads were trimmed by using the cutadapt plugin. The trimmed single-end (forward) sequences were subsequently denoised with the DADA2 plugin of QIIME2 (2). To obtain qualified data, we truncated reads to 130 bp from 3´ end based on the quality score. The DADA2 outputs high confident amplicon sequence variants (ASVs) through quality filtering, denoising and remove chimeric reads. The classify-consensus-vsearch plugin was applied for taxonomy assignment by aligning against SILVA 132 99% 16S rDNA sequences, and identity cutoff was set at ≥80% sequence similarity by default.

***Fecal short-chain fatty acid analysis***

The SCFAs we analyzed included acetic acid (C2), propionic acid (C3), butyric acid (C4), isoutyric acid (C4), valerie acid (C5), isovaleric acid (C5). Twenty mg raw feces were dissolved in 500 μL 0.5% H3PO4 aqueous solution and homogenized with Geno/Grinder® at 1,000 rpm for 2 min. Then the sample solutions were centrifuged at 18,000 × *g* for 10 min at 4 ℃ to remove the deposition. Afterwards, 285 μL upper Sample solution were collected in a new centrifuge tube, and 15 μL acetate-d3 were added. For the liquid-liquid extraction of SCFAs, 300 μL butanol and the final solution were mixed and centrifuged. Lastly, 20 μL internal standard propionate-d5 were added to 180 μL upper organic layer. GC-MS analysis was performed by Agilent 7890A gas chromatograph (Agilent Technologies, CA, USA) coupled with Pegasus 4D GC x GC-TOF MS system (Leco Corporation, USA) using an VF-WAXms capillary column (3).

***Fecal bile acid analysis***

The bile acids quantification included 4 primary bile acids ($\alpha$-MCA, $\beta$-MCA, CA, and UDCA), 4 secondary bile acids ($\omega$-MCA, CDCA, DCA, and LCA), and 7 conjugated bile acids (T$\beta$-MCA, TUDCA, TCA, GCA, TCDCA, TDCA, TLCA). Twenty milligrams raw feces were dissolved in 200 μL 70% d3-cholic acid aqueous solution containing 2 ppm d4-cholic acid as internal standard and homogenized with an ultrasonicator for 30 min. Then the sample solutions were centrifuged at 18,000 × *g* for 5 min. LC-MS analysis was performed by a Dionex U3000 UPLC system coupled with a high-resolution Q Exactive Plus instrument equipped with the ESI source (Thermo Scientific, Germany) and an Acquity HSS T3 (2.1 × 100 mm, 1.7 µm) column (Waters, USA).

***Biochemical analysis of the FGF15-FGFR4 pathway***

For western blotting, frozen tissues were lysed in lysis buffer containing 7 M urea, 2 M thiourea, 2% CHAPS, 0.002% bromophenol blue, 60 mM DTT, and a protease and phosphatase inhibitor cocktail. The cell lysates were sonicated for 5 min and centrifuged at 17,500 × *g* for 30 min at 4 °C. Total protein content was measured by Bio-Rad (Hercules, CA, USA) protein assay. For preparing the loading buffer, 62.5 mM Tris-HCl, 10% glycerol, 2% SDS, and 0.01% bromophenol blue were mixed with the protein samples, then heated at 95 °C for 10 min to denature the protein. Proteins were separated by 10% or 12% SDS−polyacrylamide gel electrophoresis gel, then transferred onto polyvinylidene difluoride membranes (Millipore, Burlington, MA, USA). After blocking with 5% bovine serum albumin in tris-buffered saline with Tween 20, the membranes were incubated with primary antibody at 4 °C for 16 h, and subsequently incubated with a secondary antibody. The protein expression signal was captured and quantified by BioSpectrum AC imaging system (UVP, Upland, CA, USA) and ImageJ (version 1.53; National Institutes of Health, Bethesda, MD, USA), respectively (4). Portal FGF15 levels were quantified using an ELISA kit (Mercodia, Uppsala, Sweden).
