## Supplementary figures 1-8 for "Dietary Exposure to Antibiotic Residues Facilitates Metabolic Disorder by Altering the Gut Microbiota and Bile Acid Composition"

***Corresponding authors:**

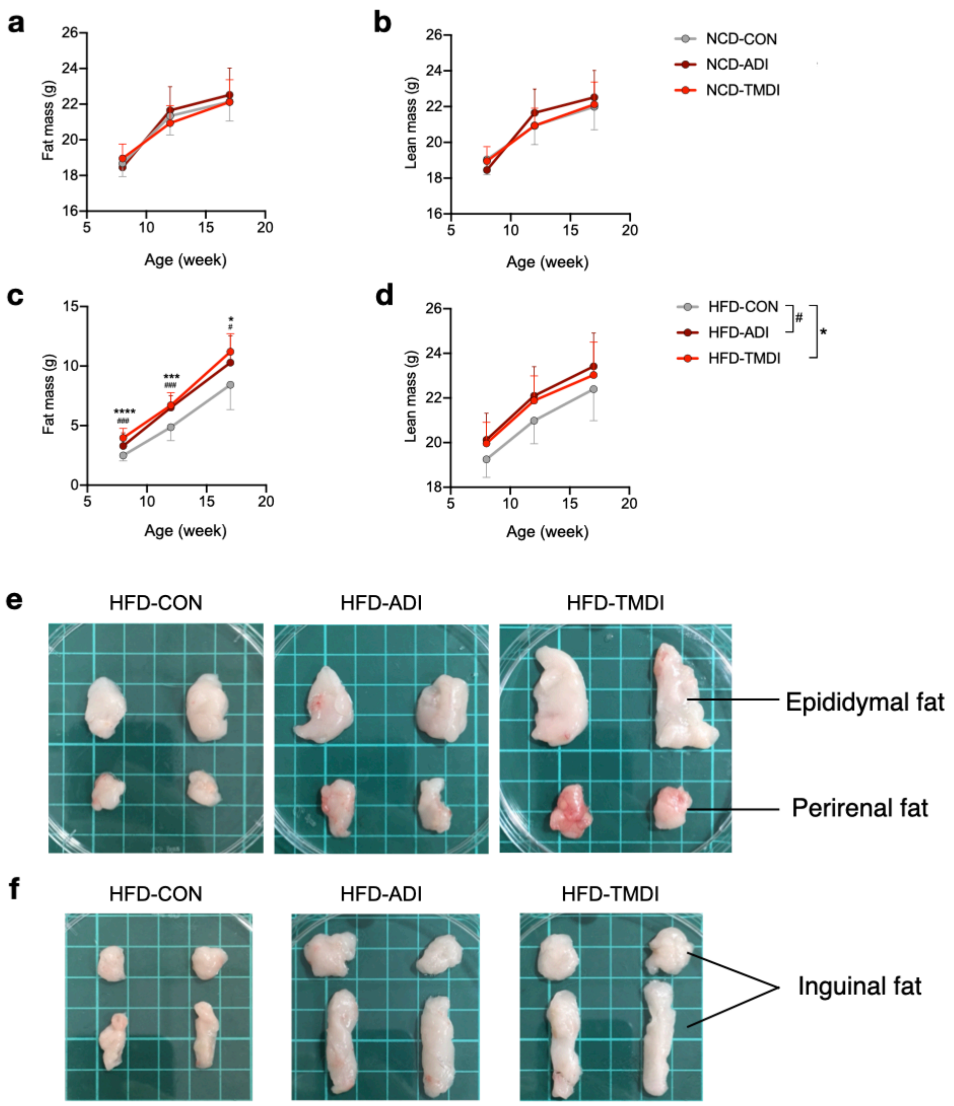


**FIG S1. Effects of tylosin on body composition and adipose tissue. (a)** Fat mass and **(b)** lean mass change of NCD mice. **(c)** Fat mass and **(d)** lean mass change of HFD mice. **(e)** Size of epididymal and perirenal fat. **(f)** Size of inguinal fat. Data are expressed as mean ± SD (n = 10–12). Statistical analyses were performed by one-way ANOVA with Tukey's range test for comparing NCD-CON versus NCD-ADI and HFD-CON versus HFD-ADI ($^{\#}p < 0.05$ and $^{\#\#}p < 0.01$), and NCD-CON versus NCD-TMDI and HFD-CON versus HFD-TMDI ($^{*}p < 0.05$; $^{**}p < 0.01$; and $^{***}p < 0.001$). Abbreviations: ADI, acceptable daily intake; CON, control; HFD, high-fat diet; NCD, normal chow diet; TMDI, theoretical maximum daily intake.

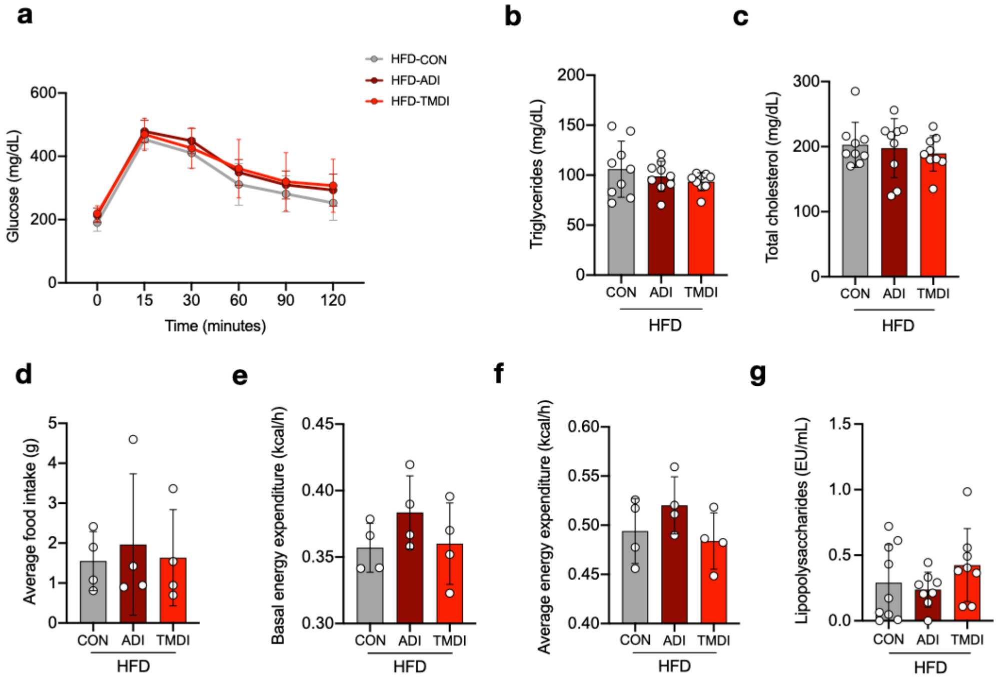


**FIG S2. Effects of tylosin on metabolic parameters. (a)** Plasma glucose profile measured during the OGTT. Plasma **(b)** triglycerides and total **(c)** cholesterol level after overnight fasting. **(d)** Average food intake. **(e)** Basal energy expenditure. **(f)** Average energy expenditure. **(g)** Plasma lipopolysaccharides level. Data are expressed as mean ± SD (n = 8–10 in OGTT and biochemistry analyses; n = 4 in metabolic measurements). Statistical analyses were performed by one-way ANOVA with Tukey’s range test for comparing HFD-CON versus HFD-ADI and HFD-CON versus HFD-TMDI. Abbreviations: ADI, acceptable daily intake; CON, control; HFD, high-fat diet; TMDI, theoretical maximum daily intake.

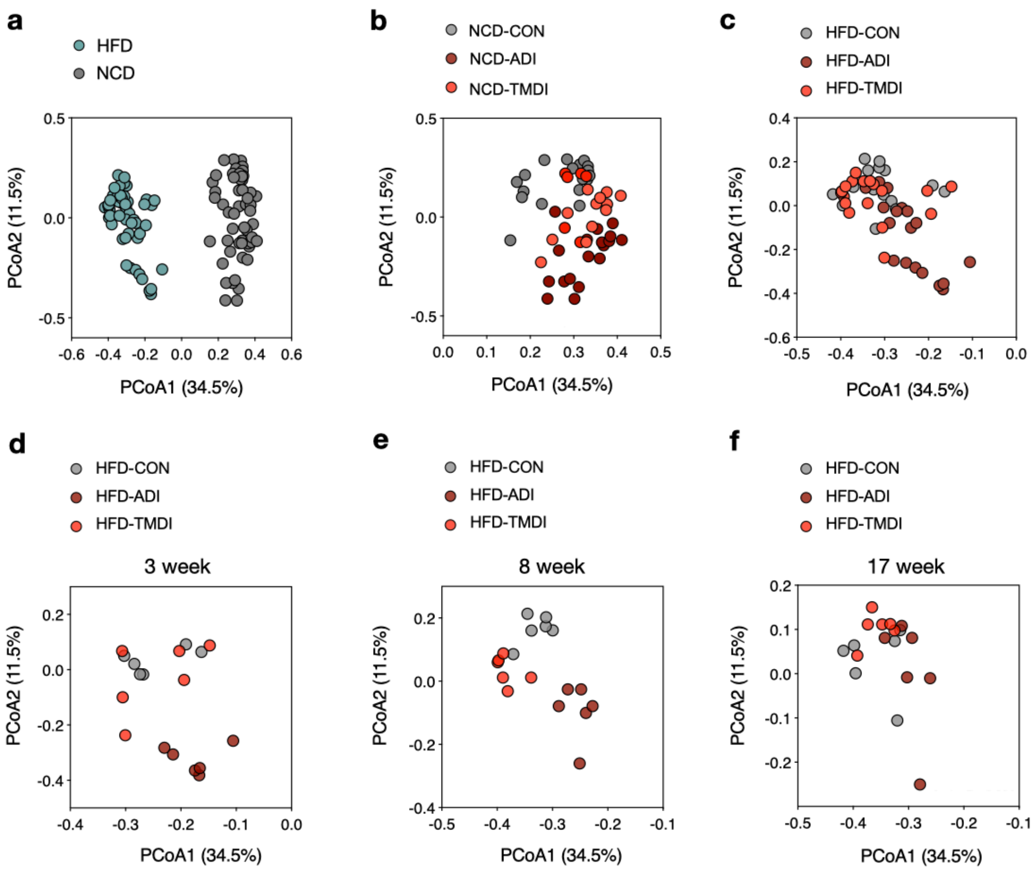


**FIG S3. Effects of tylosin on the composition of gut microbiota at different ages under different dietary conditions.** Bacterial community represented by principal coordinate analysis (PCoA) of Bray-Curtis distances of: **(a)** NCD and HFD mice; **(b)** NCD mice with different doses of tylosin; **(c)** HFD with different doses of tylosin; HFD mice at weeks **(d)** 3, **(e)** 8, and **(f)** 17 of age (n = 6). Abbreviations: ADI, acceptable daily intake; CON, control; HFD, high-fat diet; NCD, normal chow diet; TMDI, theoretical maximum daily intake.

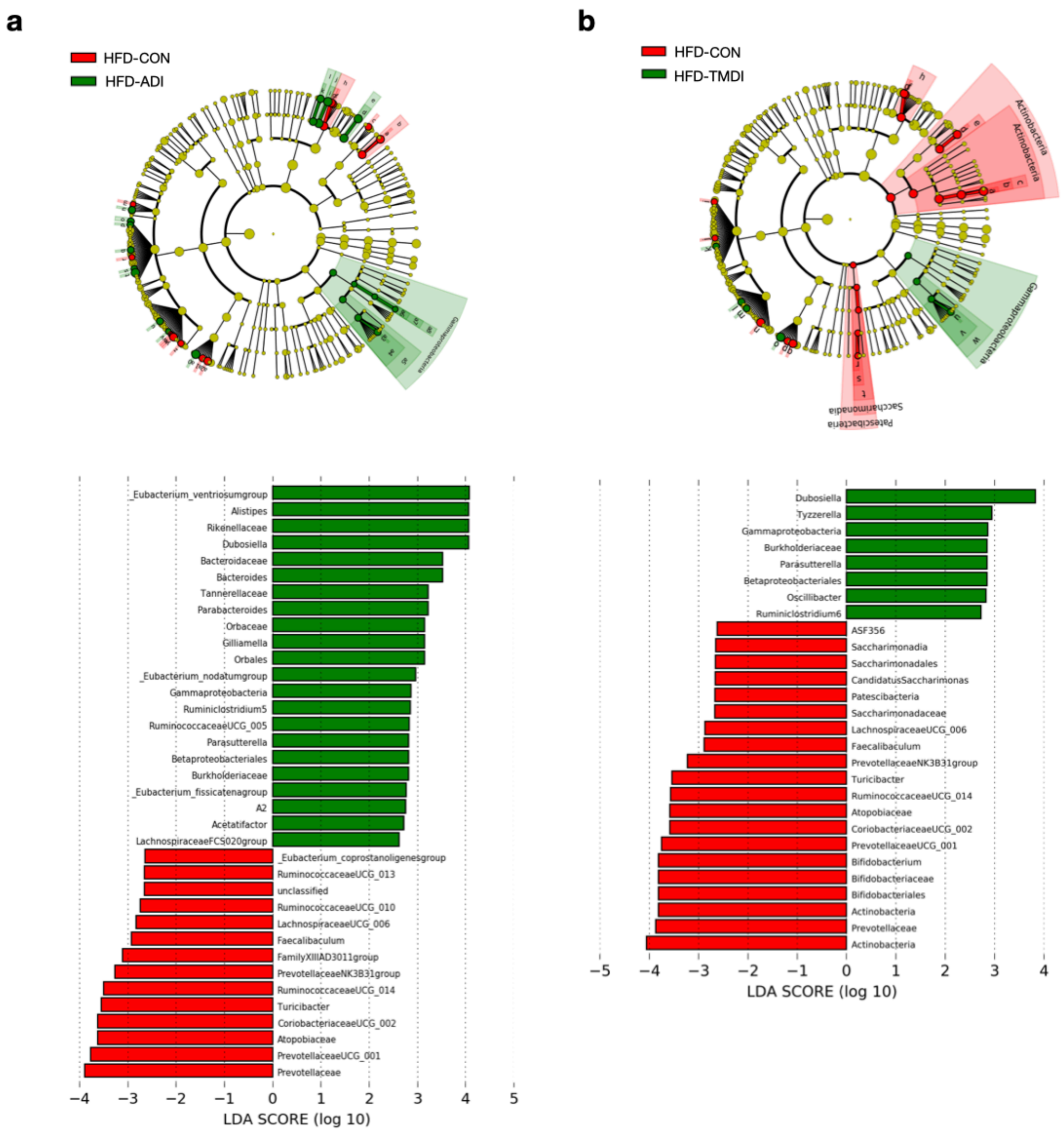


**FIG S4. Enriched bacteria** phyla **based on linear discriminant analysis effect size (LEfSe). (a)** Comparison of HFD-CON and HFD-ADI, and **(b)** HFD-CON and HFD-TMDI (n = 6). Abbreviations: ADI, acceptable daily intake; CON, control; HFD, high-fat diet; TMDI, theoretical maximum daily intake.

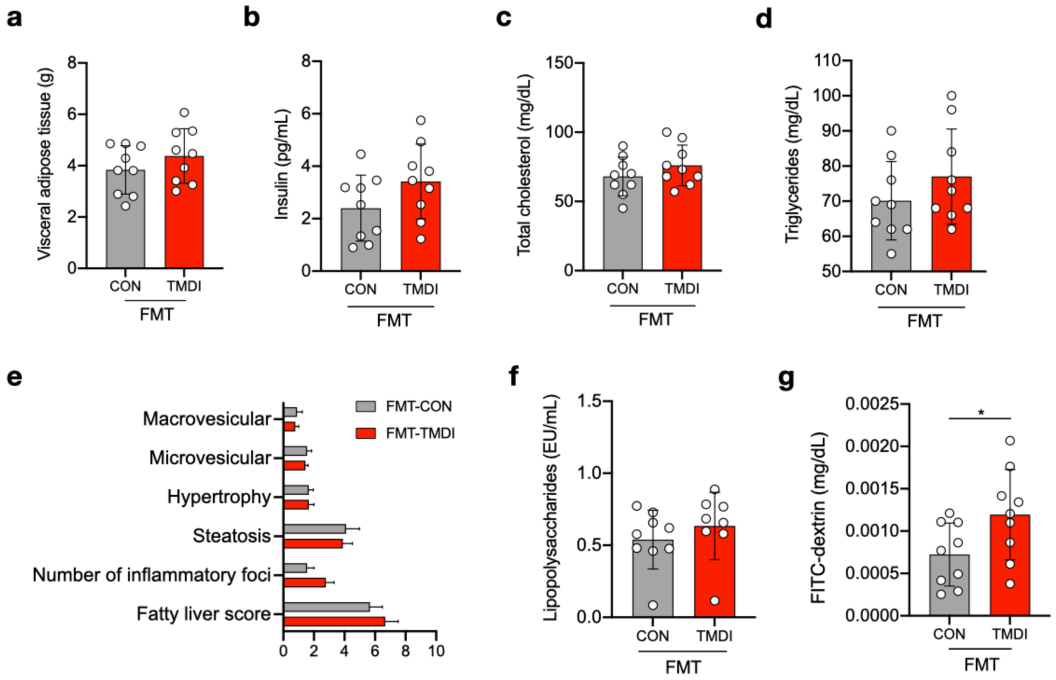


**FIG S5. Effects of transferring TMDI-tylosin-altered microbiota on metabolic parameters. (a)** Weight of visceral adipose tissue. **(b)** Plasma insulin. **(c)** Total cholesterol and **(d)** triglycerides levels after overnight fasting. **(e)** Fatty liver score. **(f)** Plasma lipopolysaccharides level. **(g)** Plasma FITC-dextrin concentration representing the intestinal permeability. Data are expressed as mean ± SD (n = 8–10). Statistical analyses were performed by unpaired *t*-test (*p < 0.05). Abbreviations: CON, control; FITC, fluorescein isothiocyanate FMT, fecal microbiota transplantation; TMDI, theoretical maximum daily intake.

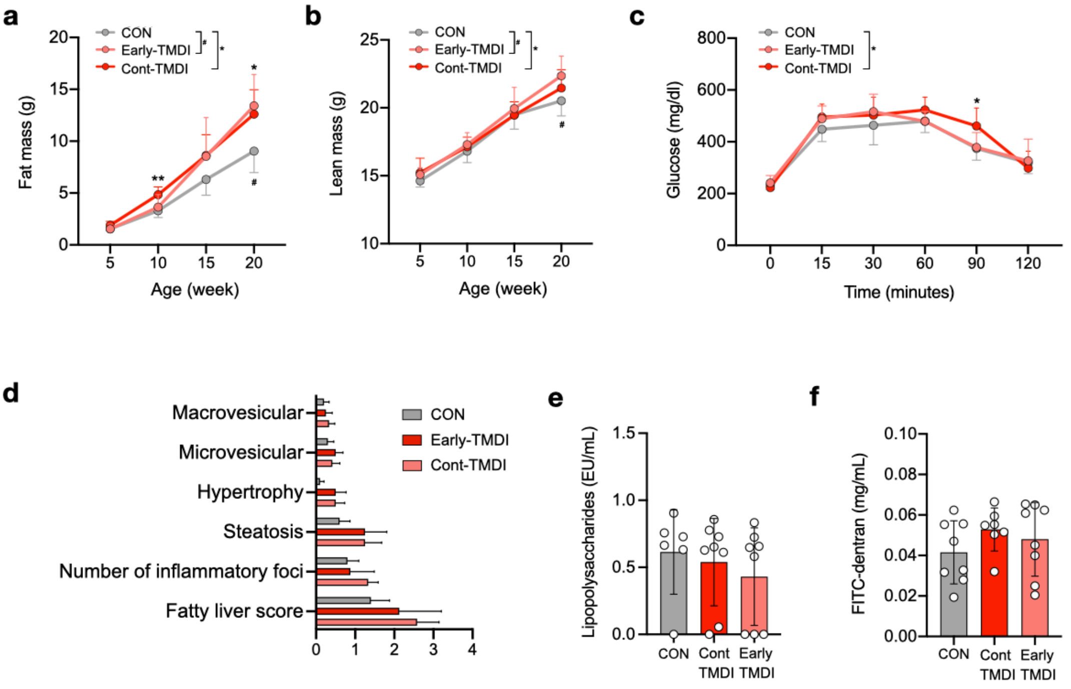


**FIG S6. Effects of early-life or continuous exposure of TMDI of tylosin on body composition, metabolic consequences, and plasma endotoxin level. (a)** Fat mass change. **(b)** Lean mass change. **(c)** AUC derived from the OGTT. **(d)** Fatty liver score. **(e)** Plasma lipopolysaccharides level. **(f)** Plasma FITC-dextrin concentration. Data are expressed as mean ± SD (n = 6–10). Statistical analysis of **(a)** and **(b)** were performed by one-way ANOVA with Tukey's range test comparing CON versus TMDI-Early ($^{\#}p < 0.05$) or CON versus TMDI ($^{*}p < 0.05$; and $^{**}p < 0.01$) at each time point. Statistical analyses of **(d–f)** were performed by one-way ANOVA with Tukey's range test. Abbreviations: CON, control; FITC, fluorescein isothiocyanate; TMDI, theoretical maximum daily intake. Treatment regimen: Cont-TMDI, continuous exposure to TMDI dose of tylosin; Early-TMDI, exposure to TMDI dose of tylosin early in life.

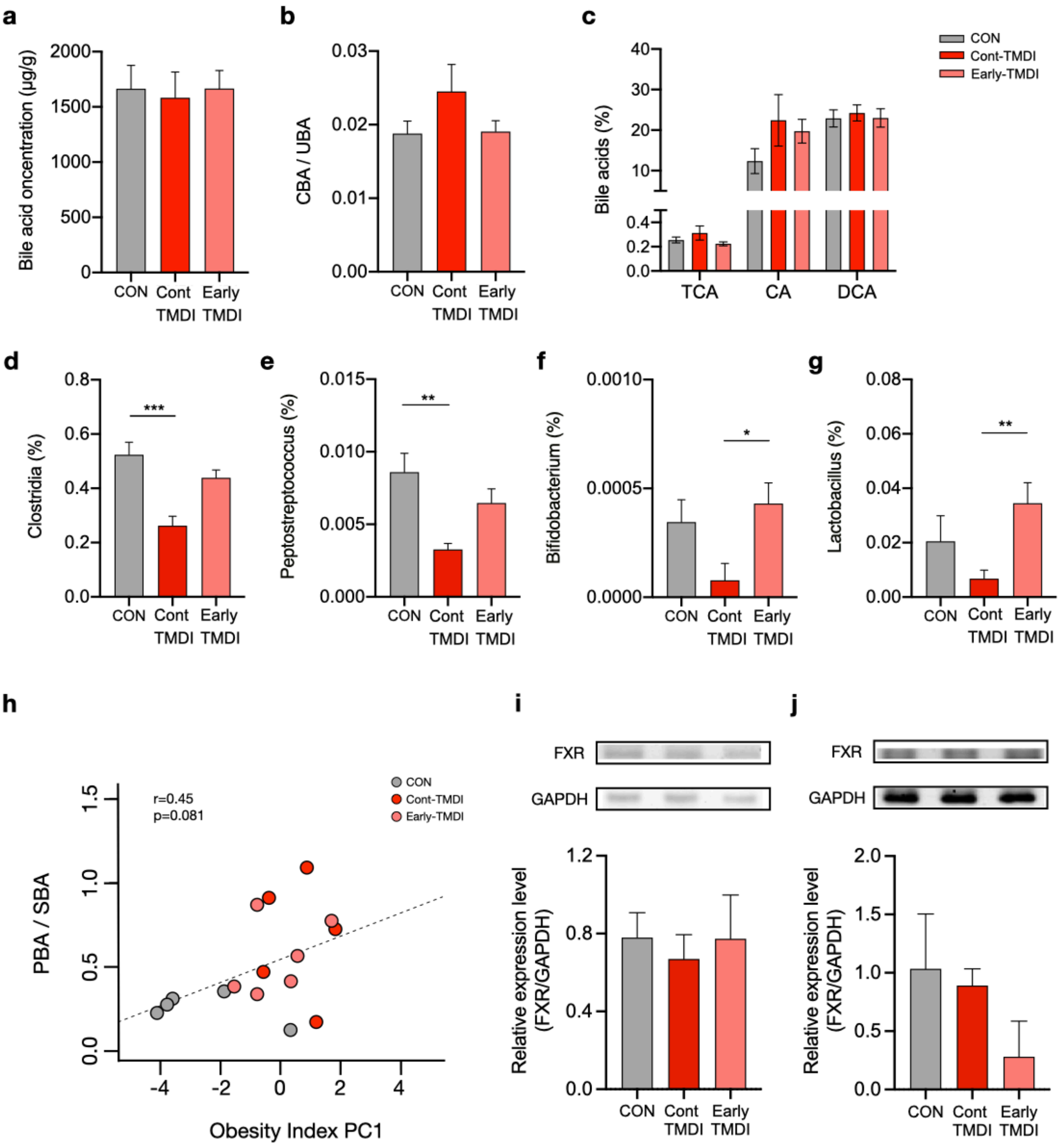


**FIG S7. Effects of tylosin TMDI on bile acid metabolism, and association between the bile acid composition and obesity.** **(a)** Total bile acid level in cecum. **(b)** Ratio of CBA to UBA; **(c)** Levels of 12-OH bile acids; **(d–g)** relative abundance of bacteria exhibiting BSH, 7α-dehydroxylase, and C-7 epimerase activity, including **(d)** *Clostridia*, **(e)** *Peptostreptococcus*, **(f)** *Bifidobacterium*, and **(g)** *Lactobacillus*. **(h)** Spearman's correlation of the ratio of PBA to SBA and the obesity index. **(i, j)** western blotting of **(i)** ileal and **(j)** hepatic FXR expression normalized to GAPDH. Data are presented by mean ± SEM in **(a–g)** and mean ± SD in **(i)** and **(j)**. For bile acids analysis, n = 8–10; for relative abundance analysis, n = 8; for western blotting, n = 4. Statistical analyses were performed by one-way ANOVA with Tukey's range test (*$p < 0.05$; **$p < 0.01$; and ***$p < 0.001$). Abbreviations: CA; cholic acid; CBA, conjugated bile acid; CON, control; DCA, deoxycholic acid; FXR, farnesoid X receptor; GAPDH, glyceraldehyde 3-phosphate dehydrogenase; PBA, primary bile acid; SBA, secondary bile acid; TCA, taurocholic acid; TMDI, theoretical maximum daily intake; UBA, unconjugated bile acid. Treatment regimen: Cont-TMDI, continuous exposure to TMDI dose of tylosin; Early-TMDI, exposure to TMDI dose of tylosin early in life.
